# Auditory representations of words during silent visual reading

**DOI:** 10.64898/2025.12.12.693932

**Authors:** Jiawei Li, Adrien Doerig, Radoslaw Martin Cichy

## Abstract

Silent visual reading is accompanied by the phenomenological experience of an inner voice. However, the temporal dynamics and functional role of the underlying neural representations remain unclear. Here, we recorded electroencephalography (EEG) data while humans read naturalistic narratives, and applied computational modelling to isolate time-resolved auditory from visual and semantic representations. Our results revealed robust auditory representations during silent reading that were not explained by visual or semantic features, emerging already before word onset. These auditory representations mimicked the sequence of sounds of the corresponding words in a fine-grained manner, revealing a candidate basis for the phenomenological experience of hearing an inner voice while reading. Finally, we show that auditory word representations exhibit a key signature of predictive processing: they are stronger for unexpected than expected words. More specifically, early auditory features contribute to predictions before word onset, whereas later features only contribute after word onset, suggesting distinct prediction stages. Together our results reveal the temporal dynamics and functional role of auditory representations in silent reading.

## Introduction

Language understanding in the human brain involves complex computations that support communication, learning, and knowledge sharing^1–7^. Modelling this complex process computationally has been a long standing challenge in neuroscience^8–15^ for both of its canonical forms: listening and reading. While a large number of computational modeling studies of language processing have focused on listening^8,9,16–19^, comparatively few empirical studies addressed reading^10,20–25^. In doing so, these studies viewed reading as involving two distinct stages: visual processing, followed by semantic processing^26–28^, assessing typically one of the two. Accordingly, computational models of reading focus on either the visual encoding of text^24,29,30^, or the semantic representations that capture its meaning^10,25,31^.

This computational modelling approach of reading is fundamentally incomplete. A large body of empirical research has shown that reading is not restricted to visuo-semantic computations, but is an integrated audio-visuo-semantic process: during reading, not only visual, but also corresponding auditory representations of words emerge. For example, early behavioral experiments showed that auditory representations accelerate or alter the course of visual word processing^32,33^. Further, reading ability is highly correlated with auditory processing abilities for basic sounds, as well as prosodic and phonological features in both healthy and dyslexic populations^34–37^. Neurally, fMRI studies showed that auditory cortex co-activates during reading tasks^3,38–42^, and EEG studies have provided evidence for the processing of auditory and semantic representations^22,43,44^. This has led to dual-route theories of reading, positing that both visual and auditory representations are co-activated during reading tasks^45–49^. These theories are also phenomenologically supported, as humans commonly report experiencing an inner voice while reading^40,50–52^.

Here, we establish a modelling approach to account for the integrated audio-visuo-semantic nature of reading. We reveal and characterize the neural dynamics that underlie auditory representations during reading on a word-by-word level. To constrain complex modelling with rich data, and to capture language processing in its natural complexity and context dependency, we employ an experimental strategy that employs complex naturalistic narratives^17,53–55^.

To anticipate, the results of our computational modelling establish the existence of uniquely auditory representations - not predicted by visual or semantic factors - during reading of naturalistic narratives, as predicted by psychological dual-route theories. On this basis we characterized auditory representations during silent reading in two ways: they are tightly temporally organized, encoding the temporal sequence of corresponding word sounds^56–58^, and they serve a predictive function as indexed by stronger effects for unexpected versus expected words^9,59–63^.

## Results and Discussion

To assess brain responses during reading of naturalistic narratives, we recorded an electroencephalography (EEG) dataset from 10 participants. Each participant silently read ∼3 hours of naturalistic narratives (**Fig. 1a**). Words were presented one by one, timed such that each word was presented with the same onset and duration as in the audio stream from which the written version was derived^19^. These ∼30 written Mandarin narratives totalled ∼25,000 words, providing a strong empirical basis for building encoding models to determine the time-resolved contribution of visual, auditory and semantic features in capturing word-specific brain dynamics^21,22,64^ during silent reading.

**Figure 1.**
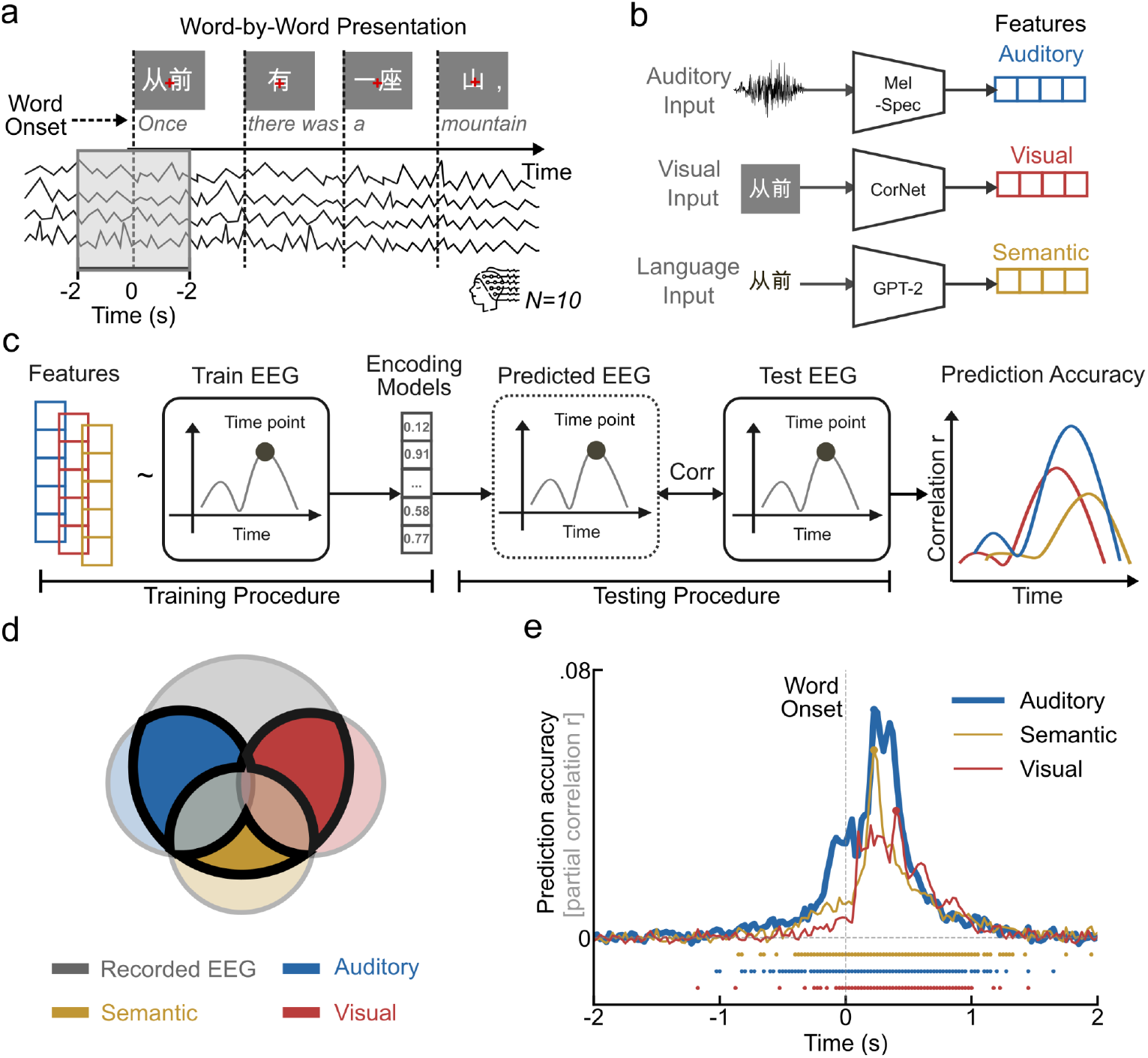
Experimental paradigm, encoding analysis pipeline, and results for modelling the neural dynamics of auditory, visual and semantic representations. **a)** Experiment paradigm. Participants were presented with complex narratives (4-7 minute length news reviews covering a range of topics such as technology and culture) word by word timed to source auditory recordings. **b)** For each word, we extracted auditory features from a Mel spectrogram, visual features from CORnet, and semantic features from GPT-2. **c)** We trained encoding models to predict EEG responses (epoched form -2s to +2s) with respect to word onset, using auditory, visual, and semantic features separately. We used the trained encoding models to predict EEG responses to held-out test words, and compared (Pearson’s *r*) predictions to the corresponding empirical EEG responses, resulting in prediction accuracy time courses. **d)** We used partial correlations to determine the unique contribution from each feature set (i.e. auditory, semantic and visual) in predicting recorded EEG (grey). Light colors indicate total contribution related to each feature set, dark colors the unique contribution of each feature set in predicting EEG data when controlling for the two other feature sets respectively. **e)** Prediction accuracy time courses indicating unique contributions of auditory, visual and semantic features in predicting EEG. Prediction accuracies are averaged across all participants and EEG channels. Dashed vertical line indicates word onset. Rows of asterisks at the bottom of the plots indicate significant time points (one-sided *t*-test, *p* < 0.05, FDR corrected across 161 time points, *N* = 10 participants).

### The temporal dynamics of auditory, visual, and semantic representations in silent reading

To reveal the time courses with which visual, auditory and semantic representations of words emerge during reading, we built encoding models using three different feature sets (**Fig. 1b**): i) auditory features derived from a Mel-frequency spectrogram^65,66^; ii) low-level visual features using responses from the first layer of a visual Deep Neural Network (CORnet) trained to simultaneously classify natural images and images of single written words^24,29^; and iii) semantic features, using the embeddings of the large language model (LLM) GPT-2^9,11,19^. We trained individual encoding models to predict the activity of each EEG electrode at each timestep (every 25 ms) of an EEG epoch from -2 to +2 s with respect to word onset using 10-fold cross-validation (**Fig. 1c**).

This procedure yielded time courses of raw prediction accuracy for each model (**Supplementary Fig. 1)**. However, because the feature sets are expected to be correlated to each other in naturalistic settings, the raw time courses do not reflect the specific contribution of each feature set. To quantify the unique contribution, we performed a partial correlation analysis (**Fig. 1d**). We measured the correlation between the predicted brain activity by an encoding (e.g., auditory) and the target EEG pattern, while controlling for the contributions of the predictions from other encoding models (e.g., visual and semantic). We assessed significance of participant-averaged prediction accuracy at each time using one-sided *t*-tests against 0 (*p* <.05, FDR-corrected for 161 time points; *N* = 10 participants).

Our results indicated that all three encoding models uniquely predicted neural responses during silent reading of words in naturalistic narratives (**Fig. 1e**, for peak latencies with 95% confidence intervals see **Supplementary Table 1**).

Concerning visual representations, this result is expected, given that visual representations in the brain are well predicted by artificial deep neural networks^24,29,67–71^. This results reveals the time course of visual processing of low-level word features, and serves as a health check for our analysis approach.

Concerning semantic representations, our results reveal the time courses with which semantic representations emerge for reading. These are qualitatively equivalent to time courses found for listening^9,59,63^, thus supporting theories positing of a common semantic access for reading and listening^10,61,72,73^. Notably, semantic processing of a word during reading emerged before the word’s onset, as reported previously for reading across ECoG^9,59^ and MEG^59,63^. This adds to the idea that our brain and LLM models share computational principles.

Concerning auditory representations, our results support dual-route theories of reading^44,45,48^ by directly assessing neural signals during silent reading. This complements previous studies that supported dual-route theories indirectly by demonstrating a modulatory function of independent auditory information on behavior or brain responses to reading. Specifically, our study provides evidence that auditory processing is involved in reading in naturalistic, complex settings^53–55^, with ecological validity that goes beyond studies that relied on more constrained and artificial paradigms, such as single word or sentence tasks^32,33,40,43,47^. Our results are also broadly consistent with previous fMRI studies reporting that auditory regions, such as superior and middle temporal gyrus (STG, MTG)^3,38–41^, co-activate when people read short words or sentences. This invites fMRI studies using encoding models and silent reading of naturalistic narratives to probe the spatial locus of the identified auditory representations, with the STG/MTG as a prediction.

### Auditory representations in silent reading represent fine-grained sequences of word sounds

Given that auditory representations of word sounds are evoked during silent reading, what kind of information do they represent? We assess two hypotheses about their temporal structure, inspired by previous work in speech processing^58,74–76^. Inspired by observations for complex auditory textures^74,77^, the first hypothesis is that the brain represents a summary statistic, i.e. a static average format of the sound without temporal structure. The alternative hypothesis stipulates that auditory representations evoked during silent reading track the fine-grained temporal sequence of word sounds, akin to the sequence emerging during speech processing ^56–58^.

To arbitrate, we determined whether there is a temporal correspondence between the sequence of auditory features of a word spoken and emerging auditory representations during reading. For this we partitioned the auditory features of the Mel spectrogram for each word into 10 discrete 20 ms time bins (i.e., the first time bin covers 0–20 ms, the second 20–40 ms, etc.) from 0–200 ms with respect to word onset (**Fig. 2a**). Then, we trained an encoding model using the model features of each bin individually to predict EEG responses with respect to word onset.

**Figure 2.**
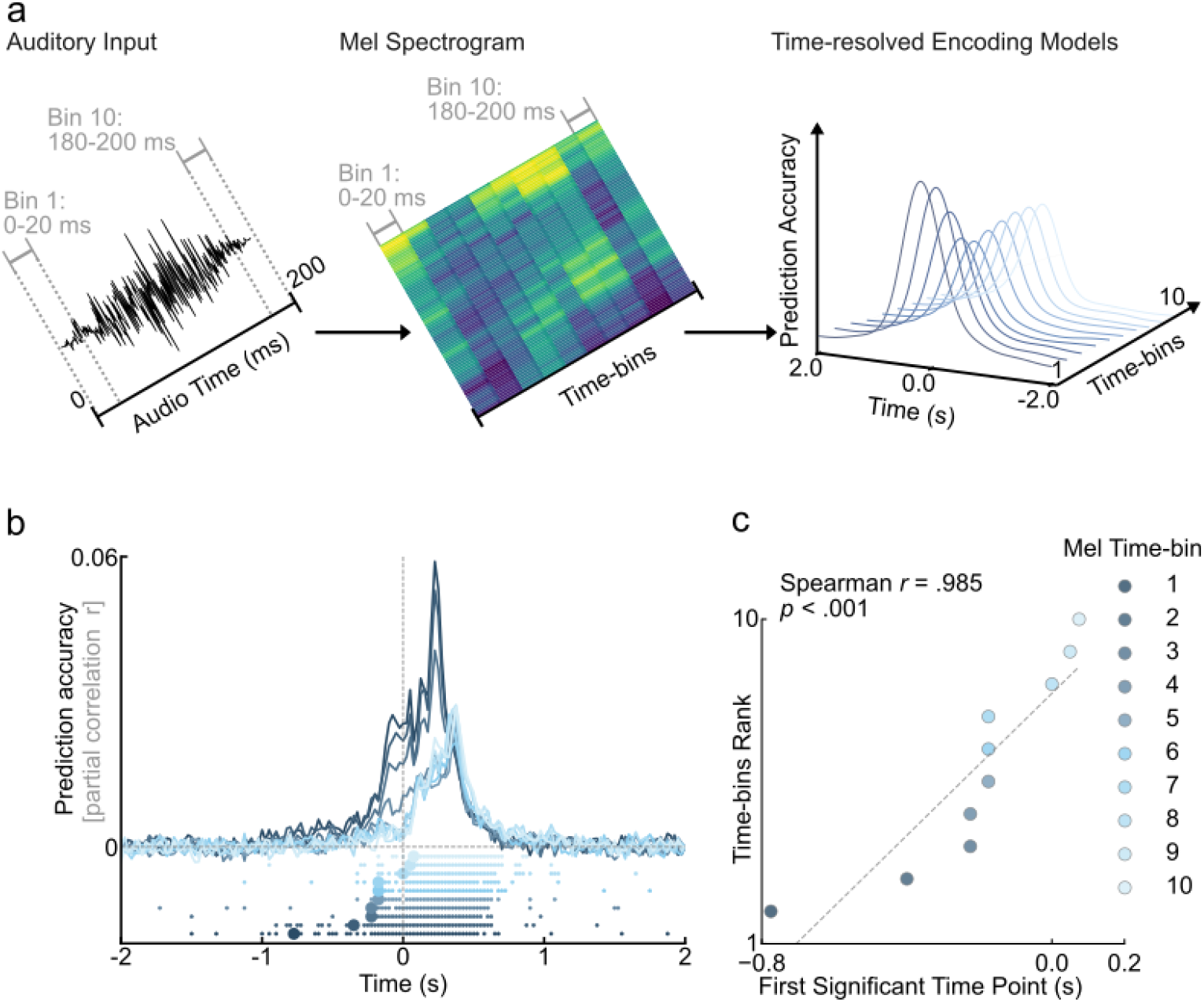
Time-resolved auditory encoding models: procedure and results. **a)** We partitioned the auditory features of the Mel-frequency spectrogram for each word into 10 discrete 20 ms time bins. We trained and evaluated encoding models based on features for each bin, yielding 10 prediction accuracy time courses. **b)** Prediction accuracy time courses for each bin (later ∼ brighter blue). Small dots indicate significance (one-sided *t*-test, *p* < 0.05, FDR-corrected across 161 time points, *N* = 10 participants), large dots mark the first time point with > 3 consecutive significant time points. Dashed vertical line indicates word onset. **c)** Scatterplot shows the relationship between Mel time-bin and significance onset, revealing strong positive correlations (Spearman *r*(8) =.985, *p* <.001).

After partialling out the visual and semantic predictions, we found that auditory features from all bins predicted EEG responses during silent reading (**Fig. 2b**), with a clear temporal order: earlier auditory features encoded earlier EEG responses, and later auditory features encoded later responses. We directly quantified this observation by calculating the Spearman correlation between the earliest time point at which auditory representations significantly predicted the neural response for >3 consecutive timepoints (**Fig. 2b** large dots) and the time-bin rank. We found a strong and significant effect (*r*(8) =.985, *p* < .001, **Fig. 2c**), demonstrating that the brain represents auditory features of words during silent reading with a fine-grained temporal sequence resembling that of the spoken word.

This finding impacts our understanding of silent reading in several ways. First, it supports dual-route theories^44–48^ of silent reading, detailing the format of the auditory representations evoked in silent reading as fine-grained and sequential. This contrasts with ideas that logographic languages such as Chinese, that is languages for which characters map onto syllables, evoke auditory representations less strongly than alphabetic languages for which single letters map onto sounds^78,79^. The auditory representations automatically emerging here are much finer-grained than the duration of a typical syllable of ∼150–250 ms, thus demonstrating the emergence of auditory information at a finer level than directly encoded logographically.

Second, it provides a putative basis for the phenomenological experience of an inner voice during silent reading. In that the auditory representations follow a temporal sequence of the corresponding sounds, the experience of the inner voice is indeed akin to veridical listening, rather than the illusion of a detailed percept^80^.

Third, relatedly, the presence of a temporal sequence suggests a functional role for these representations. Previous work in speech processing showed that the brain preserves temporal precision in auditory representations when tasks demand detailed operations on auditory stimuli^58,76,81,82^, otherwise only a summary statistic is maintained^74^. Inferring in reverse, the detailed temporal structure of auditory representations during listening supports the idea of a functional role for auditory representations in silent reading.

### Early and late auditory representations in silent reading reflect distinct prediction processes

Why does the brain adopt fine-grained auditory representations in silent reading? Prediction plays an important role during language processing in both reading and writing, involving all known representational stages from sensory to semantic representations^9,16,16,60–62,83^. We thus hypothesized that the newly identified auditory representations emerging during silent reading are related to predictive processing.

We further specified this hypothesis in two ways, guided by theoretical reasoning and the observed fine-grained temporal structure of auditory representations. First, previous literature suggests that there are two important stages for prediction: before word onset, when the prediction is formed; and after word onset, when word processing is impacted by predictability, for example related to prediction error processing^62,83–85^. We thus assessed neural responses separately before (-0.3 to 0 s)^63,84–86^ versus after word onset (0.2 to 0.5 s)^9,60–62^ separately. We selected these time windows based on prior prediction studies: The -0.3 to 0 s window captures pre-onset prediction formation^63,84–86^, while the 0.2 to 0.5 s window reflects word processing modulated by these predictions^9,60–62^. Second, previous literature suggests that early versus late auditory representations of words emerging during listening play different roles in prediction^33,56,87^. Speakers emphasize the beginnings of words^88,89^, early word representations are processed more accurately^90,91^, and consequently speech perception models emphasize early word representations^90,92^. We thus assessed early versus late emerging auditory representations of a word separately.

Classical studies reported larger stronger brain responses for semantically unexpected than expected words^9,59–63^. Analogously, we tested if the accuracy of our auditory encoding model is higher for unexpected than for expected words. For each word in our dataset, we classified it as “expected” given its context if it was among the top-5 words predicted by an LLM, and as “unexpected” if it was not in the top-5 (**Fig. 3a**). Using the LLM’s word output probability we verified that the unexpected words have higher surprisal value (i.e. lower probability assigned by the LLM) than the expected words (**Supplementary Fig. 2**). We then trained auditory encoding models separately for expected versus unexpected words, for each of the 10 different Mel spectrogram timebins as above, and partialled out the predictions of visual and semantic encoding models as before (**Supplementary Fig. 3** for each Mel spectrogram timebin). We then aggregated results as to whether they pertain to early (0–100 ms after word onset, **Fig. 3b,c**) versus late (i.e., 100–200 ms, **Fig. 3d,e**) auditory word features.

**Figure 3.**
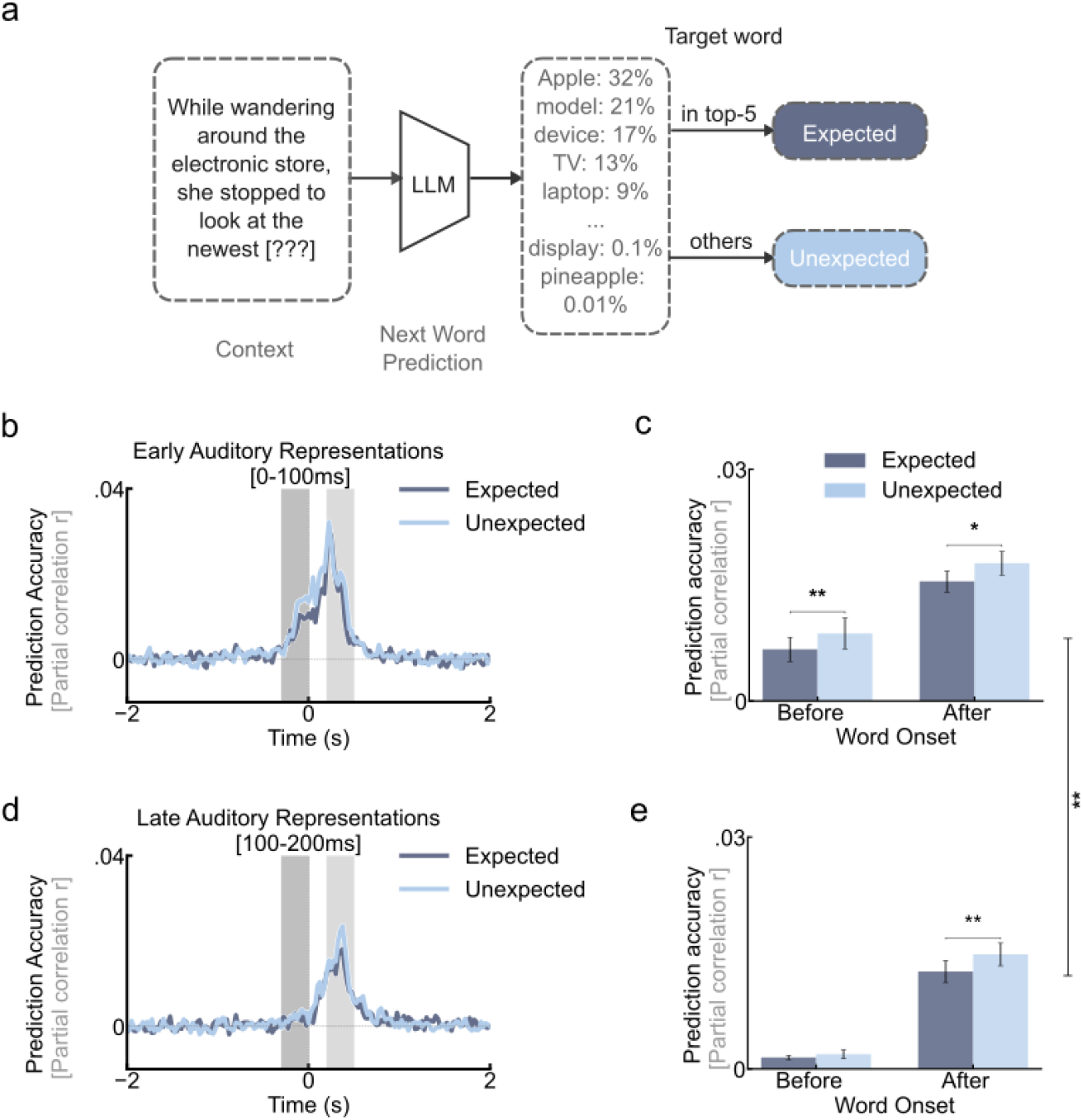
Early and late auditory representations reflect distinct prediction processes. **a)** We split words into expected vs. unexpected according to the next word predictions of LLM: if the next word fell within the LLM’s top-5 predictions, it was labeled expected (dark blue); otherwise unexpected (light blue). (**b,d**) Prediction accuracy time courses for early (b) and late (d) auditory representations. Dark shaded area indicates time window of interest before word onset (-0.3 to 0 s), light shaded area indicates time window of interest after word onset (0.2 to 0.5 s). (**c,e**) Difference in prediction accuracy between expected and unexpected words for early (c) and late (e) auditory representations (* *p* < .05, ** *p* < .01, FDR-corrected).

To assess our results statistically, we conducted a three-way repeated-measures ANOVA with the following factors: time of representation (i.e., early vs. late auditory features), word onset relationship (i.e., predicting EEG before vs. after word onset), and word expectancy (i.e. training encoding models on expected vs. unexpected words). This yielded significant main and interaction effects (**Supplementary Table 2**).

The main effect of word expectancy (*F*(1,9) = 17.31, *p* =.002, **Fig. 3c,e**), with higher prediction accuracy for unexpected compared to expected words, indicating a role of auditory representations in predictive processing. The main effect of time of representations (*F*(1,9) = 16.79, *p* =.003), with higher prediction accuracy for early versus late auditory representations, highlights the importance of early auditory representations in prediction. This dichotomy in processing early versus late auditory representations of words observed here in silent reading mirrors the dichotomy during listening^56,58^, suggesting common mechanisms. The main effect of relationship to word onset (*F*(1,9) = 63.24, *p* < .001, **Fig. 3c,e**), with higher prediction accuracy after than before word onset, suggests differing processes at those two stages. Interpreted from the standpoint of predictive coding^93–95^, those stages would map on prediction proper before word onset, and prediction error processing after word onset.

Among the interaction effects, we observed a significant three-way interaction (*F*(1, 9) = 7.86, *p* =.021) and a two-way interaction of word onset relationship × time of representation (*F*(1, 9) = 5.79, *p* =.040). Investigation with post-hoc tests revealed that before onset, unexpected words elicited significantly higher prediction accuracy than expected words for the early auditory representations (*t*(9) = 3.41, *p* =.011, FDR-corrected for all the following reported *p*-values), but not for the late auditory representations (*t*(9) = 0.71, *p* =.495). In contrast, after word onset, unexpected words elicited significantly higher prediction accuracy than predictable words for both the early representations (*t*(9) = 3.20, *p* =.014) and the late representations (*t*(9) = 4.03, *p* =.005). Our interpretation is that when predicting a word before its onset, the brain generates only the beginning of the auditory representation rather than the full form as an efficient means to guide further processing. After word onset, however, the brain processes the entire auditory representation of the word, for better informed processing guidance beyond the first guess before word onset.

## Summary

Collectively, our results reveal the characteristics of fine-grained auditory representations in silent reading of complex and naturalistic narratives. This fleshes out the tenets of the dual-route theories of reading^45,46^ in neural terms, and argues for the ecological validity of those theories. The fine-grained temporal and sequential nature of auditory representations in reading offer a suitable basis for the phenomenological experience of an inner voice while reading. Auditory representations finally show hallmarks of predictive processing, indicating their functional role in language understanding.

## Materials and Methods

### Participants

Ten healthy participants (seven females; mean age = 26.56 ± 3.75 years) completed six EEG sessions to acquire neural data during reading to short Chinese narratives. All participants had normal or corrected-to-normal vision, and were native Chinese speakers. All participants provided informed written consent and received monetary reimbursement. Procedures were approved by the ethics committee of the Department of Education and Psychology at Freie Universität Berlin.

This sample size is based on earlier studies using encoding models to model human language processing^19,25,73^ that also used an intensive sampling strategy, extensively recording individual participant data in order to ensure sufficient training and testing data for within-participant encoding models^19,54,96,97^.

### Experimental stimuli and paradigm

The stimulus set was extracted from a publicly available dataset^19^ consisting of audio recordings and written, human-checked transcripts of 60 news reviews, each lasting 4 to 7 minutes, covering a varied range of topics such as technology and culture. The narratives range in duration from 240 to 406 seconds. They contain in total 43,340 words, with each narrative comprising 516 to 906 words, together using a vocabulary of 10,492 unique words. The mean word length was 0.33 seconds, with the first quartile at 0.13 seconds, the third quartile at 0.57 seconds, and a standard deviation (SD) of 0.18 seconds (**Supplementary Fig. 4**).

We presented participants with sequences of single words in white font on a gray background, overlaid with a central red fixation cross^73,98,99^. The duration of each word matched the length of the corresponding word in the source audio recordings. Depending on word length, the visual angle of the visual presentation ranged from 2×2° (single Chinese character) to 2×10° (five Chinese characters). Punctuation was attached to the previous characters^73,98,99^(**Fig. 1a**).

Ten participants took part in multiple EEG sessions of 2h each. Participant 1 read 51 narratives (43,923 words), participant 2 read 36 narratives (33,665 words), participant 3 read 24 narratives (22,188 words), and participants 4-10 read 30 unique narratives (ca. 25,000 words). For each participant narratives were randomly selected from the 60 available narratives in the full stimulus dataset^19^ with the exception of a single narrative that was presented in every session and for every participant (not analyzed here to avoid repetition effects^100^). To prevent order effects, the presentation order of the narratives was randomized for every participant. Reading narratives was further interspersed with other tasks that are not included in the analysis here.

### EEG recording and preprocessing

We recorded the EEG data using a 64-channel EASYCAP with electrodes arranged in the standard 10–10 system, and a Brainvision actiCHamp amplifier at a sampling rate of 1,000 Hz. We removed blink and muscle artifacts using independent component analysis (ICA), re-referenced the data to the average reference, downsampled it to 200 Hz, and filtered the data to 1–40 Hz using the MNE toolbox in Python. We epoched the EEG trials from -2 s to 2 s with respect to word onset^9,66^ (**Fig. 1a**) as the basis for encoding model training and testing.

### Feature extraction

As feature basis sets for building encoding models we extracted auditory, visual and semantic features from different computational models. Common to all feature sets, after extraction, we applied principal component analysis (PCA) to reduce dimensionality to 50 and equate dimensionality across models for fair comparison.

#### Auditory features

To extract auditory features from the recorded narratives, we used Mel-frequency spectrograms. In detail, for each word, we extracted the 0–200 ms window after onset (10 timebins, each 20 ms). To obtain a summary auditory feature set, we flattened the resulting 80 × 10 timebins into an 800-dimensional acoustic feature vector, in line with previous work^66^. To obtain time-resolved auditory features, we kept the 10 discrete 20 ms timebins (i.e., the first time-bin covers 0–20 ms, the second 20–40 ms, etc.).

#### Visual features

To extract visual features, we extracted activations from images of words (**Fig. 1b**) using a CORNet model trained specifically on a combination of natural images and written words^24,29^. We flatten the extracted activations for subsequent analysis. We used the first layer (i.e. model layer-V1) activations of the model.

#### Semantic features

To capture contextual semantic information, we used embeddings provided by the authors of the dataset containing our stimuli^19^. These embeddings were extracted using a GPT-2 model with a context window size of 1024 tokens. The model has 25 layers^19^ and each layer’s embedding is a 1024-dimensional vector for the target word. Guided by previous literature we chose layer 12 to predict brain activity^8,18^.

### Encoding models

We trained encoding models using linear regression to predict brain responses from the embeddings described in *Feature extraction*^9,70^. We used a 10-fold cross-validation approach for training, using Pearson correlation between the predicted and true brain activities as the performance measure (**Fig. 1c**).

We trained encoding models separately for each timepoint, (i.e., every 25 ms within the time window of -2 s to 2 s with respect to word onset^9^). This analysis was carried out for each EEG channel separately. We trained models separately based on auditory, visual, and semantic features (**Fig. 1c**).

### Partial correlations to determine unique feature contribution

To assess the unique contribution of a feature set, we computed the partial correlations between the recorded EEG signals and the EEG signals predicted from those features (e.g. auditory features), while controlling for the contributions of the EEG predicted from the other two features (e.g., visual and semantic features) (**Fig. 1d**).

### Expected vs. unexpected words

Words were categorized as either expected or unexpected based on predictions from the open-access Chinese BLOOM-1B4 model (https://huggingface.co/Langboat/bloom-1b4-zh). This model is different from the GPT-2 model used in our encoding models. This was required by the fact that this version of GPT-2 is not open access^19^. We evaluated each word in context, using all preceding words as input to generate a prediction for the target word. If the word was in the top five words predicted by the model, it was labeled as an “expected” word; otherwise, it was labeled as an “unexpected” word. This resulted in 16,732 expected words (38.61%) and 26,608 unexpected words (61.39%) in the whole dataset. For each word, the surprisal value was defined as − *log P(Word|context)*, with probabilities derived from the language model’s next-word prediction distribution.

We trained encoding models separately for expected vs. unexpected words, and compared their prediction accuracy. To ensure equal sample sizes for training and testing, we randomly subsampled the set of unexpected words to match in number the set of expected words.

### Pre-word onset and post-word onset time windows

To assess neural responses separately before versus after word onset in terms of prediction processes, we chose time windows based on previous literature: The -0.3 to 0 s window captures pre-onset prediction formation^63,84–86^, while the 0.2 to 0.5 s window reflects word processing modulated by these predictions^9,60–62^.

### Statistical testing

To establish the statistical significance of the encoding model results, we tested all results against 0 using one-sided *t*-tests. To correct for multiple comparisons, we applied Benjamini-Hochberg false discovery rate (FDR) correction^101^ to the resulting *p*-values to account for the number of EEG time points (*N* = 161) in all analyses.

To calculate 95% confidence intervals for both the peak latencies and the differences in peak latencies, we generated 1,000 bootstrapped samples by resampling the participant-specific results with replacement. This approach produced empirical distributions of the respective measures, from which we derived the confidence intervals.

We conducted a three-way repeated-measures ANOVA with the factors time of representation (early vs. late auditory features), word onset relationship (before vs. after word onset), and word expectancy (expected vs. unexpected words). The ANOVA assessed main effects and interactions among these factors. We performed post-hoc paired *t*-tests to explore simple main effects within each factor combination, and *p*-values were corrected for multiple comparisons using FDR-correction.

## Data and code availability

The dataset and the code will be openly available upon publication.

## Acknowledgments

This work was supported by a German Research Council (DFG) grant (INST 272/297-2) (R.M.C.), a European Research Council (ERC) Consolidator grant (ERC-CoG-2024101123101) (R.M.C.), and a Humboldt Scholarship (J.L.). We thank the HPC Service of FUB-IT, Freie Universität Berlin, for computing time (https://doi.org/10.17169/refubium-26754).

