## Supplementary Fig. 1 for "Auditory representations of words during silent visual reading"

**Supplementary Figure 1** | Raw prediction accuracy time courses for each feature set

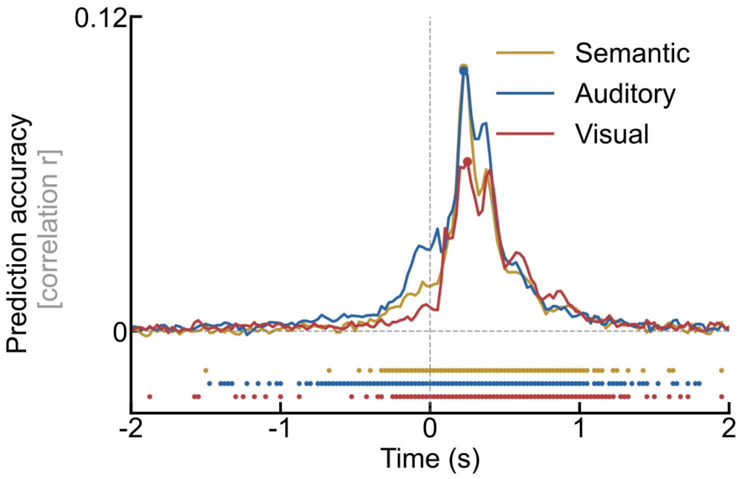


The prediction accuracies are averaged across all participants and EEG channels. The dashed vertical lines indicate the onset of word. Rows of asterisks at the bottom of the plots indicate significant time points (one-sided *t*-test, $p < 0.05$, FDR corrected across 161 time points, $N = 10$ participants).
