## Supplementary Fig. 2 for "Auditory representations of words during silent visual reading"

**Supplementary Figure 2** | The distribution of the surprisal values of the expected vs. unexpected words

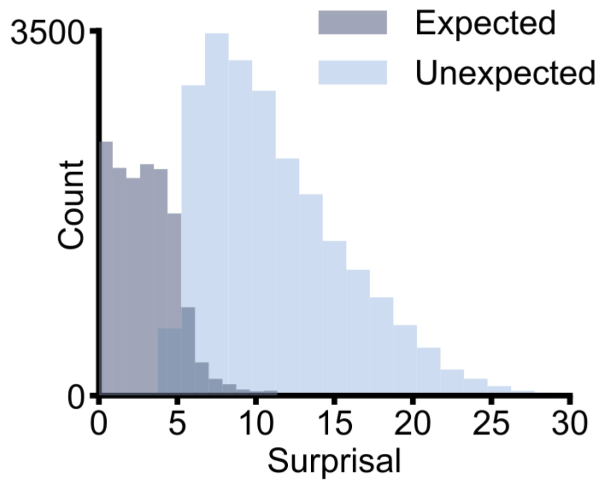
