## Supplementary Fig. 3 for "Auditory representations of words during silent visual reading"

**Supplementary Figure 3** | Time courses of prediction accuracy of each mel-spectrogram time bins for expected vs. unexpected words

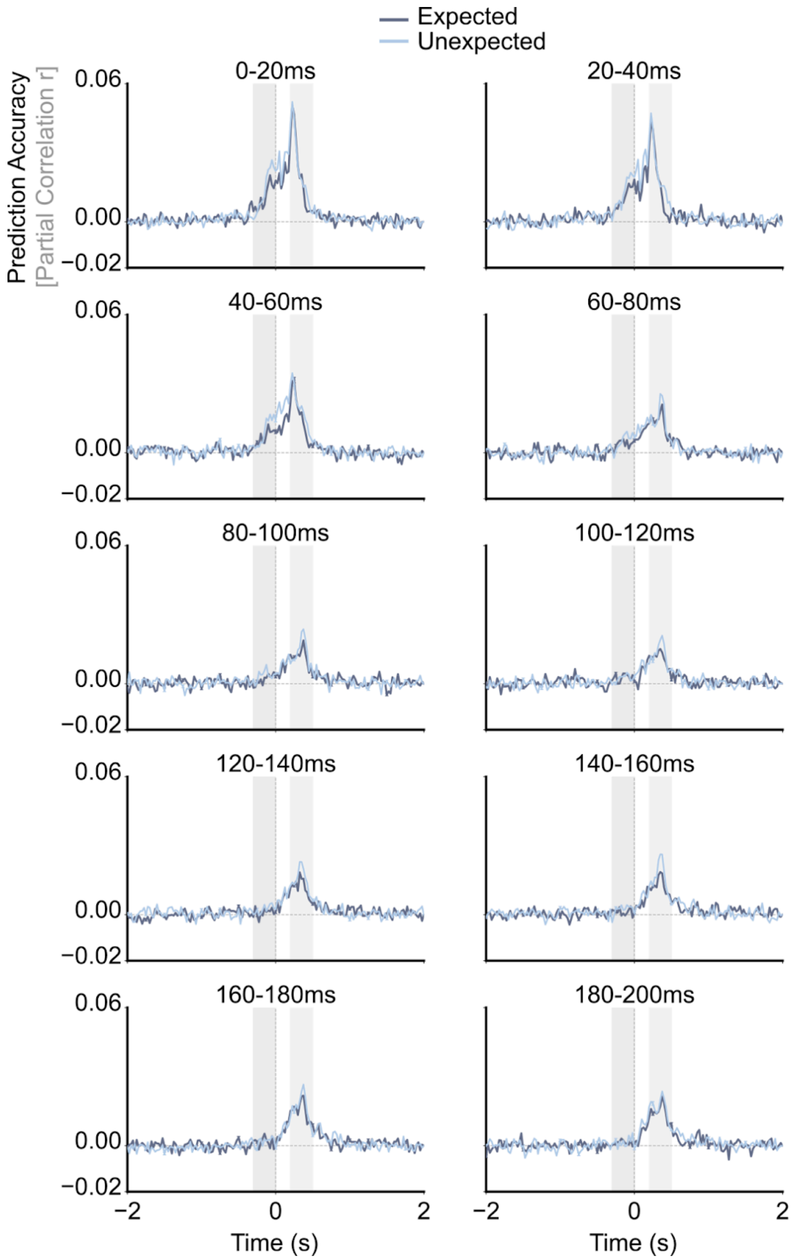


The dark blue lines show the results for the expected words, and the light blue lines show the results for the unexpected words. The prediction accuracies are averaged across all participants and EEG channels. The dashed vertical lines indicate the onset of word. The dark grey shadow indicated the early time window(-0.3 to 0s), and the light grey shadow indicated the late time window(0.2 to 0.5s).
