## Supplementary Fig. 4 for "Auditory representations of words during silent visual reading"

**Supplementary Figure 4** | Distribution of word lengths across all stories

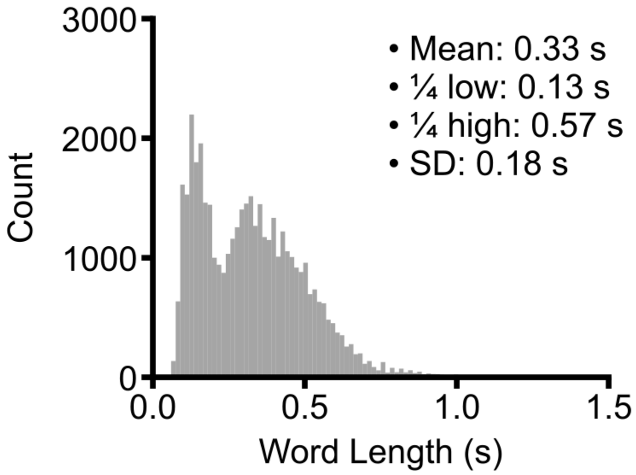


The mean word length was 0.33 s, with the first quartile(1/4) at 0.13 s, the third quartile at 0.57 s, and a standard deviation(SD) of 0.18 s.
