## Supplementary Table 1 for "Auditory representations of words during silent visual reading"

**Supplementary Table 1** | Bootstrapped peak latency of time courses showing unique contributions for the three models

| Model | Peak Latency (ms) | 95% CI (ms) | Peak Prediction Accuracy |
|---|---|---|---|
| Auditory | 252 | [225, 350] | 0.068 |
| Semantic | 227 | [225–250] | 0.056 |
| Visual | 379 | [200–425] | 0.038 |
