## Supplementary Table 2 for "Auditory representations of words during silent visual reading"

**Supplementary Table 2** | Word Onset × Word Expectancy × Time of Representation Anova Statistical Details

| | *F*(1,9) Value | *p* Value |
|---|---|---|
| Word Onset | 63.24 | < .001 |
| Word Expectancy | 17.31 | .002 |
| Time of Representation | 16.79 | .003 |
| Word Onset × Word Expectancy | 1.86 | .206 |
| Word Onset Relationship × Time of Representation | 5.79 | .040 |
| Word Expectancy × Time of Representation | 4.10 | .074 |
| Word Onset × Word Expectancy × Time of Representation | 7.86 | .021 |
